# Androgen Receptor promotes renal cell carcinoma (RCC) vasculogenic mimicry (VM) *via* altering TWIST1 nonsense-mediated decay through lncRNA-TANAR

**DOI:** 10.1101/2020.06.30.180067

**Authors:** Bosen You, Yin Sun, Qing Liu, Keliang Wang, Ruizhe Fang, Bingmei Liu, Fuju Chou, Jie Luo, Ronghao Wang, Jialin Meng, Chi-Ping Huang, Shuyuan Yeh, Wanhai Xu, Chawnshang Chang

**Affiliations:** Department of Urology, The 4th Affiliated Hospital of Harbin Medical University, Harbin 150001, China; Department of Urology, The 2nd Affiliated Hospital of Harbin Medical University, Harbin 150001, China; George Whipple Lab for Cancer Research, Departments of Pathology and Urology, and The Wilmot Cancer Institute, University of Rochester Medical Center, Rochester, NY, USA, 14646; Department of Pathology and Cutaneous Oncology, Heilongjiang Provincial Hospital, Harbin 150001, China; Sex Hormone Research Center and Departments of Urology, China Medical University/Hospital, Taichung 404, Taiwan

**Author notes:** These authors contributed equally to this work (Bosen You, Yin Sun, Qing Liu, Keliang Wang). Correspondence to: Wanhai Xu and Chawnshang Chang.

**Keywords:** AR, Vasculogenic mimicry, lncRNA, Nonsense-mediated mRNA decay

## Abstract

While the androgen receptor (AR) may influence the progression of clear cell renal cell carcinoma (ccRCC), its role to impact vasculogenic mimicry (VM) to alter the ccRCC progression and metastasis remains obscure. Here we demonstrated that elevated AR expression was positively correlated with tumor-originated vasculogenesis in ccRCC patients. Consistently, *in vitro* research revealed AR promoted VM formation in ccRCC cell lines *via* modulating lncRNA-TANAR/TWIST1 signals. Mechanism dissection showed that AR could increase lncRNA-TANAR (TANAR) expression through binding to the androgen response elements (AREs) located on its promoter region. Moreover, we found that TANAR could impede nonsense-mediated mRNA decay (NMD) of TWIST1 mRNA by direct interaction with TWIST1 5’UTR. A preclinical study using *in vivo* mouse model with orthotopic xenografts of ccRCC cells further confirmed the *in vitro* data. Together, these results illustrated that AR-mediated lnc-TANAR signals might play a crucial role in ccRCC VM formation and metastasis, and targeting this newly identified AR/lncRNA-TANAR/TWIST1 signaling may help in the development of a novel anti-angiogenesis therapy to better suppress the ccRCC progression.

## Introduction

Clear cell renal cell carcinoma (ccRCC), the major subtype of aggressive human malignancies, accounted for approximately 1,000,000 cases and 175,000 deaths worldwide in 2018 (1). Despite the shift towards early stages at diagnosis, fatal metastasis or rapid recurrence occurred in 1/3 to 1/4 of cases (2). For most metastatic RCC, anti-angiogenesis therapy, such as treatment with sunitinib and pazopanib, have proved its efficacy as well as Vascular Endothelial Growth Factor Receptor (VEGFR)-Tyrosine Kinase Inhibitor (TKI). However, the vast majority of patients will ultimately acquire resistance and relapse, and some patients are inherently refractory to the targeted therapy (3, 4). Hence, the detailed mechanisms of RCC progression and angiogenesis need to be comprehensively understood for the development of better efficacies.

The androgen receptor (AR) can promote progression and hematogenous metastasis in ccRCC through ASS1P3/miR-34a-5p/ASS1 and miR-185-5p/HIF-2a/VEGF signaling, respectively. This is consistent with the higher incidence and more malignant phenotypes in males according to data from the Surveillance Epidemiology and End Results (SEER) database (5–7). Furthermore, targeting AR with the antiandrogen enzalutamide could restore sunitinib sensitivity in the sunitinib-resistant PDX mouse model, suggesting enhancing TKI efficacy through inhibition of AR to better suppress ccRCC progression (8). Paradoxically, a higher expression of AR is associated with better prognosis based on a retrospective analysis from the TCGA database, indicating that AR function in ccRCC is complex and may be tumor stage-dependent (9).

Besides the classical tumor angiogenesis, Maniotis *et al*. demonstrated a *de novo* pattern of tumor perfusion, named vasculogenic mimicry (VM), which is directly surrounded by cancer cells (10). Due to its ability to circulate blood from vessels to tumor tissues and to facilitate tumor cells into the extracellular matrix, VM plays a significant role in boosting tumor growth as well as promoting metastasis, which contributes to poor prognosis and aggressiveness in many cancers (11–16). On the other hand, as an alternative nutrient supplement, VM complements the cancer vasculature theory and provides a mechanistic alternative for the inherently or acquired resistance to anti-angiogenesis therapy (18). Periodic acid Schiff (PAS) staining and CD31 (or CD34) immunohistochemistry (IHC) have been widely used to identify VM formation *in vivo* (10). In addition to matrigel architecture, a novel collagen-induced migration program of VM, characterized by short fibers and small pores provides another *in vitro* assay to mimic the VM formation *in vivo* (19). Although this unique vascular channel formation was first reported in kidney cancer in 2013 (20), to date the VM molecular pathway in RCC still remains poorly understood.

Long noncoding RNAs (lncRNAs), a class of untranslated transcripts longer than 200 nucleotides barely with protein-coding capacity, participate in diverse biological processes (21). Mounting evidence has proved lncRNAs significantly influence the pathogenesis of cancers *via* transcriptional regulation or direct interaction with miRNAs, mRNAs, and proteins in a multitude of molecular pathways (22). Recently, lncRNAs have been shown to also modulate genitourinary malignancies *via* different pathways including proliferation, apoptosis, angiogenesis, and drug resistance. However, little is known about the potential role of lncRNAs involved in VM formation (4, 23–26).

In this study, we aimed to determine the biological function of AR in ccRCC VM. Mechanism dissection revealed that the lncRNA-TANAR, which is regulated by AR, might increase the oncogene TWIST1 expression through competitive binding to TWIST1 mRNA to reduce the activity of Up-frameshift protein 1 (UPF1), the core factor required for nonsense-mediated mRNA decay (27).

## Methods and Materials

### Patients and samples

A total of 51 histologically confirmed ccRCC tissue samples with 23 paired adjacent noncancerous tissues were collected between August 1st, 2014 to February 1st, 2016 from the Department of Urology, the Second Affiliated Hospital of Harbin Medical University (Harbin, China). Patients were excluded if they had been treated previously with neoadjuvant chemotherapy or TKIs. All samples collected for use in research after patients signed the Scientific Ethics Consent were fixed in 10% formalin and then embedded in paraffin. Our research was approved by the Institutional Review Board of the hospital in advance.

### Cell culture and reagents

786O, SW839, and HEK293T cell lines were purchased from the American Type Culture Collection (ATCC, Manassas, VA). Cells were cultured in DMEM media with 1% penicillin and streptomycin, containing 10% fetal bovine serum (FBS). All cells were maintained in a humidified 5%(v/v) CO_2_ incubator at 37°C. According to ATCC’s protocol, all cell lines used in the paper have been authenticated and proven to be mycoplasma and bacteria-free and were periodically re-authenticated by PCR. Enzalutamide (South Brunswick, New Jersey) was applied at 10 μM. 5α-Dihydrotestosterone (DHT) (Sigma, St Louis, MO) was used at 10 nM. Harmine was also purchased from Sigma. Actinomycin D was purchased from Cayman Chemical (Ann Arbor, MI).

### 2D Matrigel-based tube formation assay

After incubating at 4°C overnight, 50 μl Growth factor reduced Matrigel (BD Biosciences, USA) was added to the wells of 96-well plate evenly and incubated at 37 °C for 1.5 h. Subsequently, 100 μl cells were resuspended with serum-free DMEM and loaded onto the surface of Matrigel at 2 ×10^4^ cells/well. After incubation at 37 °C for 6 h, tube formation was analyzed using microscopy (Olympus, Tokyo, Japan). Tubules were quantified by ImageJ software, which was as same as in a previous study (28). Tubule lengths in each field were photographed and an average of tubule lengths in 3-5 random fields in each well were calculated.

### 3D Collagen 1-induced tube formation assay

After mixing 10x reconstitution buffer,1:1 (v/v) with cells suspended in DMEM media, with soluble rat tail type I collagen in acetic acid (Corning, Corning, NY) to the final concentration (29), and 1 M NaOH was used to normalize pH (pH 7, 10–20 μl 1 M NaOH). Then 200 μl mixture was loaded in 48-well culture plates and incubated in a humidified 5%(v/v) CO_2_ incubator at 37°C for 7 days. Then VM formation was viewed under microscopy.

### Lentivirus packaging and cell transfection

The plasmids pLKO.1-shAR, pLKO.1-shTWIST1, pLKO.1-ENST00000425110.1 (TANAR1# and TANAR2#), pLKO.1-ENST00000436510.1, pLKO.1-ENST00000471626.1, pLKO.1-ENST00000593604.1, pLKO.1-ENST00000377977.3, pWPI-AR, pWPI-TWIST1, pWPI-TANAR wild-type, and pWPI-TANAR mutant were co-transfected with package and envelope plasmids, psPAX2 and pMD2.G into HEK293T cells for 48 h following the standard calcium phosphate transfection method to produce the lentivirus particle soup, which was then collected and stored at −80°C for later infection of ccRCC cells.

### RNA extraction and quantitative real-time PCR (qRT-PCR) analysis

Total RNAs were isolated using Trizol reagent (Invitrogen, Grand Island, NY), and 2 μg of total RNA was subjected to reverse transcription using Superscript III transcriptase (Invitrogen). Real-time PCR (RT-PCR) was conducted using a Bio-Rad CFX96 system with SYBR green to determine the mRNA expression level of a gene of interest. Expression levels were normalized to the GAPDH level using the 2^-ΔΔ^Ct methods. The sequences of the primers are listed in **Table S1**.

### Western blot

Cells were washed twice with cold PBS and lysed in cell lysis buffer and equal proteins (30-50 μg) were loading, mixed, boiled, and separated on 6–12% SDS/PAGE gel, then transferred onto PVDF membranes (Millipore, Billerica, MA). We blocked the PVDF membranes for one h *via* 5% skim milk and incubated with the specific primary antibodies overnight. After incubated with HRP-conjugated secondary antibodies, the PVDF membranes were visualized using the ECL system (Thermo Fisher Scientific, Rochester, NY). Primary antibodies used in the study for western blot follow: AR (Santa Cruz, #sc-816, Paso Robles, CA), GAPDH (Santa Cruz, #sc-166574), B-actin (Santa Cruz, #sc-517582) Argonaute-2 (Ago2) (Cell Signaling Technology, #2897, Danvers, MA), IgG (Santa Cruz, #sc-2027), TWIST1 (Abclonal #A7314, Woburn, MA), VE-cadherin (Santa Cruz, #sc-9989), UPF1 (Bethyl Laboratories, #A300-38A, Montgomery, TX), and LAMC2 (Santa Cruz, #sc-25341).

### RNA Immunoprecipitation (RIP)

Native RIP was performed as previously described. We applied RIP to perform Ago2 pull-down assay and UPF1 pull-down assays. Briefly, Cells were lysed in 1 mL ice-cold RNase-free cell lysis buffer supplemented with 1 μL RNase inhibitor (M0307S, NEB; Ipswich, MA). After storage in −80°C refrigerator for about 40 mins, the mixture was centrifuged to collect the supernatant. Then we mixed the supernatant with Protein A/G beads to preclean. Next, we incubated cell lysates with Ago2 or UPF1 antibody overnight at 4°C. The Protein A/G beads were added to each tube and rotated for 1 h. The complex was washed 8-10 times by RIP buffer and the RNA extracted using Trizol (Invitrogen).Then, qPCR was performed according to the manufacturer’s protocol (30).

### The lncRNA/mRNA pull-down assay

After receiving assigned treatments, the cells were harvested in cell lysis buffer. Detecting the levels of GAPDH *via* qRT-PCR guaranteed that the input of each group had equal loading for the following procedures. The cell lysate mixtures supplemented with 1.0 μl RNase inhibitor and 500 pMole biotin-labeled anti-sense oligos against enst00000425110.1 (5’-CAG TGT CTC CAG GTG GAC CCT GTG TCT CCT-3’) or TWIST1 mRNA (5’’-AGC TTG CCA TCT TGG AGT CCA G-3’) were rotated overnight at 4°C. The cell lysis mixtures incubated with 10 μl Streptavidin Agarose beads were rotated for 1 h at 4°C and mixtures washed by RIP buffer five times. Total RNA or protein was extracted according to the manufacturer’s protocol, and qPCR analysis or Western blots were performed to detect mRNA or protein levels, respectively (31).

### Luciferase Reporter Assay

2000bp human lncRNA-TANAR(ENST00000425110.1) promoter was cloned into PGL3 basic vectors (Promega). By mutating the crucial site of AR binding site in the lncRNA-TANAR 5’ promoter to EcoRI cutting site (-GAATTC), we constructed the mutant promoter PGL3 vector. PRL-TK was used as an internal control that served as the baseline control response. 786O and SW839 cells were plated in 24-well plates and the cDNA was transfected with Lipofectamine 3000 transfection reagent (Invitrogen) according to the manufacturer’s instructions. Luciferase activity was measured 36-48 hrs after transfection by Dual-Luciferase Assay (Promega) according to the manufacturer’s manual.

### Chromatin Immunoprecipitation Assay (ChIP)

In brief, SW839 cells were cross-linked with 3% formaldehyde and lysed in lysis buffer, then, we extracted and sonicated lysates. Protein A/G beads were used to preclear the chromatin. We then added anti-AR antibody (2.0 μg) or anti-IgG antibody to the DNA-protein mixture and incubated overnight at 4°C. Specific primer sets were designed to amplify a target sequence within the human ENST00000425110.1 promoter and agarose gel electrophoresis was used to identify the PCR products. The sequences of the primers are listed in **Table S2**.

### RNA fluorescence in situ hybridization (FISH)

FISH was performed to detect the presence of lncRNA-TANAR and TWIST1 mRNA, using a designed biotin-labeled probe (VA6-16446-VC, Thermofisher Scientific, Invitrogen™). The signals of the probes were detected by ViewRNA™ ISH Cell Assay Kit (QVC0001, Thermofisher Scientific, Invitrogen™) according to the manufacturer’s instructions and previous literature (32). The images were recorded by AxioImagerZ.1 fluorescence microscope.

### Immunofluorescence

Immunofluorescence was performed with paraffin sections according to standard procedures. In brief, the slides were deparaffinized and hydrated followed by antigen retrieval. After being blocked with 10% goat serum, the slides were incubated with the primary antibody against AR (dilution 1:200) and LAMC2 (dilution 1:200) overnight at 4°C. Next, slides were labeled with two different secondary antibodies for 1 h at room temperature. After washing three times with PBS, the slides were subsequently labeled with DAPI for visualization of staining intensity.

### *In vivo* studies

All experimental procedures were performed in accordance with the National Institutes of Health Guide for the Care and Use of Laboratory Animals, conformed to the Regulations for Animal Experimentation and reviewed by the Institutional Laboratory Animal Care and Use Committee of the University of Rochester Medical Center. Male athymic BALB/c nude mice (6–8 weeks old) were obtained from the Animal Production Area of the National Cancer Institute-Fredrick Cancer Research and Development Center in Frederick, MD, USA.

### Immunohistochemistry (IHC) staining

After fixing in 4% neutral buffered paraformaldehyde for 18 h, the patient or mouse tissue samples were embedded in paraffin and sequentially cut into 4 μm slices. After deparaffinization, hydration, antigen retrieval, and blocking, these slices were incubated with corresponding primary antibodies, then incubated with biotinylated secondary antibodies (Vector Laboratories, Burlingame, CA, USA), and finally visualized by VECTASTAIN ABC peroxidase system and 3, 3’-diaminobenzidine (DAB) kit (Vector Laboratories). The percentage of positive cells was rated per high-power field (HPF) using 400× magnification as follows: sections with 1% positive cells were rated as 0, 1 to 25% positive cells as 1, 26 to 50% positive cells as 2, 51% to 75% positive cells as 3, and 76% to 100% positive cells as 4. The staining intensity was rated as follows: 1 for weak intensity, 2 for moderate intensity, and 3 for high intensity. Points for staining intensity and the percentage of positive cells were multiplied. Tumor specimens were classified into 3 groups according to overall scoring: negative expression as 0 to 1, weak expression as 2 to 4, and high expression as 6 to 12 points. Total scores were as follows: 0 to 4 (low) and 6 to 12 (high). All slides were evaluated independently by 2 pathologists without knowledge of the identity of patients and the clinical outcome.

### Statistics

Experiments were repeated independently at least 3 times with data points completed in triplicate. Results are shown as mean ± S.D. Statistical significance was determined using the Student’s *t*-test and two-way ANOVA test by SPSS 22 (IBM Corp., Armonk, NY) or GraphPad Prism 6 (GraphPad Software, Inc., La Jolla, CA). *P* values less than 0.05 were considered statistically significant (\**P* < 0.05, \*\**P* < 0.01, \*\*\**P* < 0.001).

## Results

### AR expression may influence VM presence in ccRCC patients

To clarify the clinical significance of VM on ccRCC, we collected 51 ccRCC human specimens along with their clinicopathological data and performed the PAS/CD31 double staining to detect VM **(Fig. 1A)**. The VM was significantly higher in Stage II and Stage III compared with Stage I **(Fig. 1B)**. There is no statistical difference between males and females in any TNM Stage **(SFig. 1A)**. However, the result shows the VM occurs more frequently in males than in females, when both had a high degree of malignancy (Stage II & Stage III) **(Fig. 1C)**. Interestingly, based on Higgins Renal Microarray Dataset (33) and The Cancer Genome Atlas (TCGA) DNA Copy Number Data, we found both AR mRNA and genomic DNA content increased in ccRCC **(Fig. 1D; E)**. Next, we detected the AR expression *via* IHC in 51 ccRCC samples (36 males and 15 females), then separated patients into AR-positive group and AR-negative group **(Fig. 1F)**. As shown in **Fig. 1G**, AR expression is positively correlated with higher VM formation. Moreover, immunofluorescence (IF) analysis in human clinical samples revealed that high AR expression was positively correlated and colocalized with Laminin 5γ2, a specific VM marker (34) **(Fig. 1H)**.

**Fig. 1.**
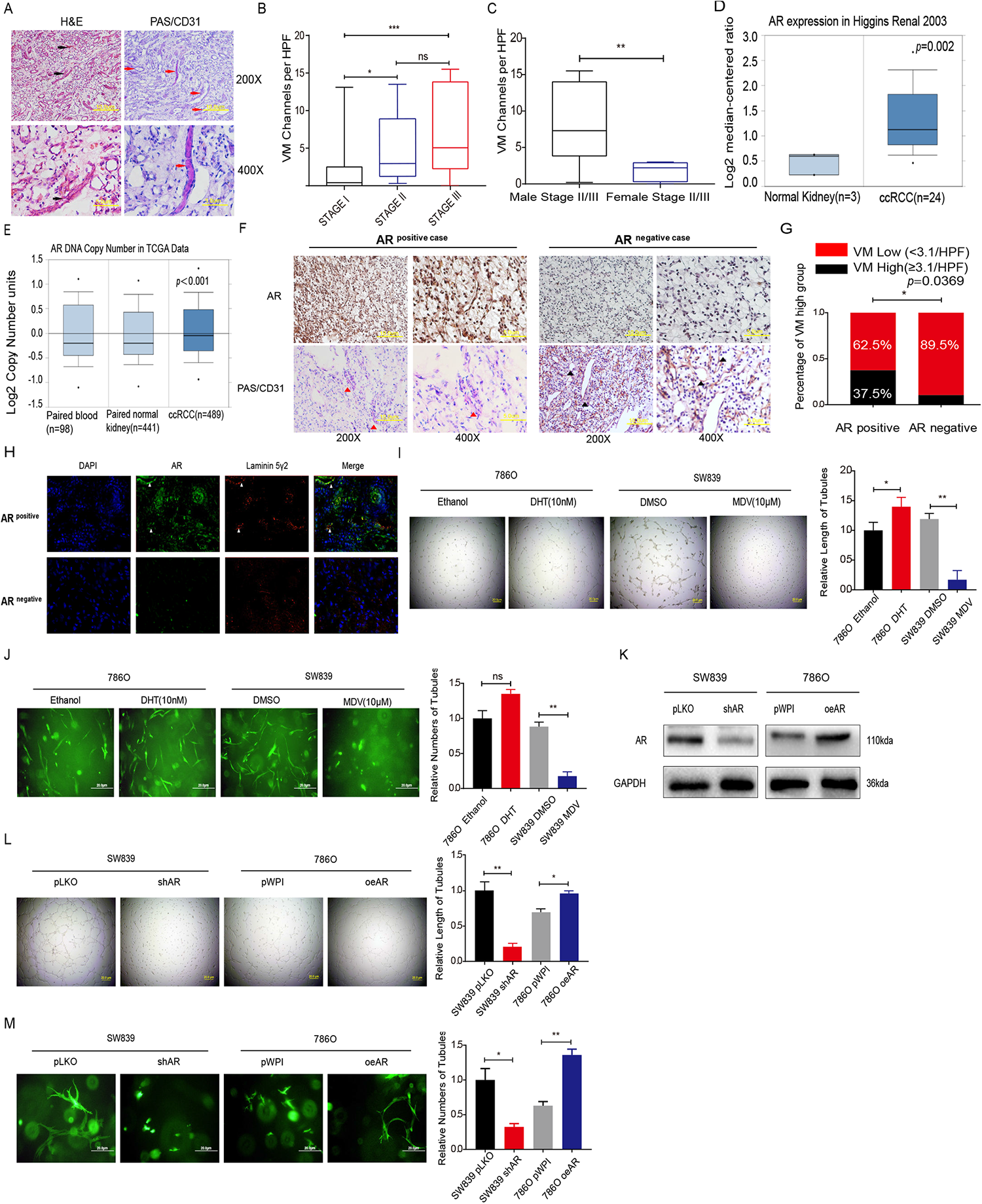
AR promotes VM formation in ccRCC. (A) CD31/PAS and HE staining were utilized to identify the VM formation. Red arrows indicate the VM channels surrounded by tumor cells with CD31 negative staining and black arrows show the VM channel with visible blood cells inside. (B) VM channels per high-quality frame (HPF) in both TNM Stage II and III are higher than in Stage I. (C) The association between gender and VM formation in ccRCC Stage II and III patients. (D) The AR mRNA expression increases in ccRCC compared with normal kidney in Higgins renal microarray dataset. (E) TCGA DNA copy number databases (TCGA, http://tcga-data.nci.nih.gov/tcga/) show higher AR DNA copy number in ccRCC than in paired normal tissue and blood. (F) VM formation per HPF quantified in AR Positive Expression Group (n=32) and AR Negative Expression Group (n=19). Red triangles show PAS+/CD31-VM channels and Black triangles show PAS+/CD31+ endothelial cell-dependent vessels. (G) Statistical analysis of the correlation between AR and VM. (H) IF analysis revealed that in AR positive sample VM numbers were more than in AR negative sample, laminin 5 gamma2 chain-positive channels were regarded as VM formation. Nuclei were stained with DAPI. (I) Matrigel-coated 2D VM tube formation assay for ethanol/10 nM DHT and DMSO/10 μM Enz treated 786O and SW839 cells, respectively. (J) 786O cells treated with ethanol/10 nM DHT (left panels) and SW839 cells treated with DMSO/10 μM Enz (right panels) were grown in Collagen I matrix for 7 days to detect 3D VM tube formation assay. (K) Western blot assay for scramble control (pLKO) and knocked down AR (shAR) in SW839 cells (left panels), vector control (pWPI) or overexpressed AR (oeAR) in 786O cells (right panels). (L-M) Matrigel-coated 2D VM assay (L) and the Collagen-based 3D VM assay (M) showed shAR in SW839 cells could suppress VM formation and oeAR in 786O cells could increase VM formation. For I, L and M, quantitations are at the right and data are expressed as mean±S.D. *p < 0.05, **p < 0.01, and ***p < 0.001 compared to the controls.

To further validate the positive correlation between AR and VM formation *in vitro*, we treated RCC 786O AR-low-positive cell line and SW839 AR-high-positive cell line with 10 nM dihydrotestosterone (DHT) and 10 μM enzalutamide (Enz), an FDA approved antiandrogen, respectively. As shown in **Fig. 1I-J**, AR agonist DHT could increase both Matrigel-coated 2D and collagen I based 3D VM formation, while AR antagonist Enz dramatically suppressed 2D & 3D VM formation only in the AR-positive cell line. Next, we knocked down the AR expression *via* adding AR-shRNA (shAR) in RCC AR-positive SW839 cells **(Fig. 1K, left panel)**, and found a decreased 2D & 3D VM formation **(Fig. 1L and M, left panel)**. In contrast, overexpressing AR *via* adding AR-cDNA (oeAR) in AR-weakly positive 786O cells **(Fig. 1K, right panel)** led to more VM formation in both 2D &3D conditions **(Fig. 1L and M, right panel)**.

Taken together, results from **Fig. 1A-M** and **SFig.1A** demonstrate that AR may play a positive role to affect the ccRCC VM formation.

### AR can induce TWIST1 to promote VM formation in ccRCC *in vitro*

To dissect the molecular basis for AR’s effect on ccRCC VM formation, we screened the known critical genes linked with VM *via* qRT-PCR assay in RCC 786O cells after adding AR-cDNA **(Fig. 2A, left panel)** or RCC SW839 cells after adding shAR plasmid **(Fig. 2A, right panel)** to examine if any of those VM-related genes may be responsive to altering AR expression (35, 36). The result showed that TWIST1 mRNA level was the only VM related gene positively correlated with AR (threshold?0.3 fold of Log10) **(Fig. 2A)**. Consistent with this, western blot assay further showed that knocking down AR in SW839 cells could decrease TWIST1, a dominant VM-promoting gene (37, 38), and overexpressing AR in 786O cells had the opposite effect **(Fig. 2B)**.

**Fig. 2.**
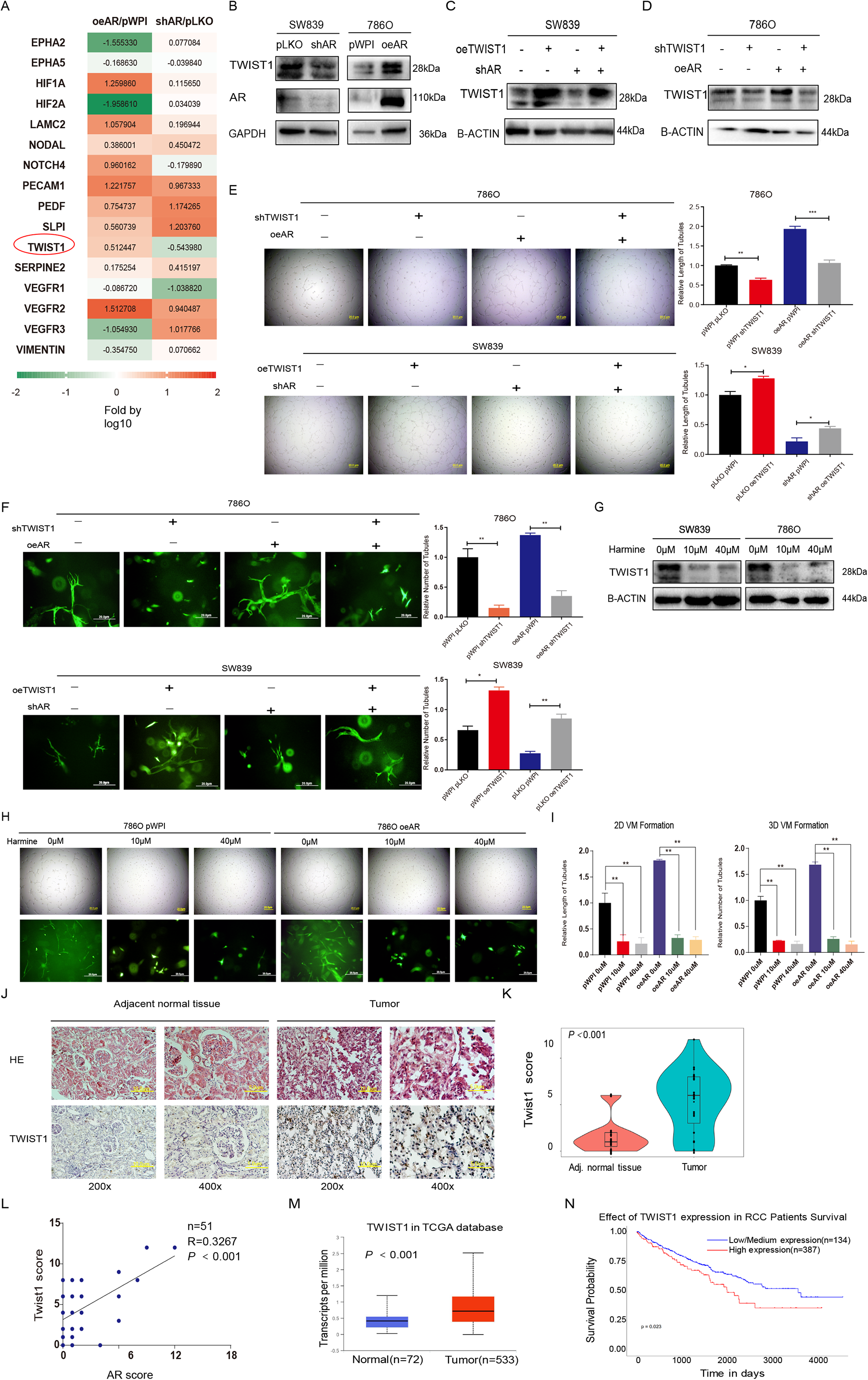
AR enhances ccRCC VM formation via altering TWIST1 expression. (A) The qRT-PCR assays for the 16 genes related to VM formation in 786O cells (left) transfected with AR-cDNA (oeAR) versus control (pWPI) and SW839 cells transfected with shRNA-AR (shAR) versus mock (pLKO). (B) Western blot assay for AR, TWIST1, and CDH5 protein levels in SW839 cells (left) with pLKO or shAR and in 786O cells (right) with pWPI or oeTR4. (C-D) Western blot assays were performed on SW839 cells (C) transfected as indicated as well as 786O cells transfected as indicated (D). (E) 2D Matrigel-coated VM assay were performed in 786O cells transfected as indicated (upper panels) and SW839 cells transfected as indicated (lower panels). (F) 3D Collagen I-based VM assay were performed in in 786O cells transfected as indicated (lower panels) and SW839 cells transfected as indicated (lower panels). (G) Western blot assays to detect TWIST1 protein levels in SW839 (left) and 786O (right) cells after treating with 10 μM or 40 μM Harmine for 48 h. (H-I) 2D Matrigel-coated (upper panels) and 3D Collagen I-based (lower panels) VM assays were performed after treating 786O cells transfected with pWPI or oeAR with TWIST1 inhibitor, Harmine. (J) Representative H&E and IHC staining of TWIST1 in adjacent noncancerous tissues (left) compared to paired ccRCC tissues (right). (K) The IHC score of TWIST1 between 23 paired adjacent tissue and ccRCC tumors. (L) The correlation between AR and TWIST1 level based on IHC score (n=51) (M) The mRNA level of TWIST in ccRCC samples (n = 534) and adjacent normal tissues (n = 72) from TCGA database. (N) Overall survival probability of ccRCC patients was negatively correlated with TWIST1 mRNA expression based on TCGA database. For E and F, quantitations are at the right and data are expressed as mean ± S.D. *p < 0.05 and **p < 0.01 compared to controls.

In order to confirm whether AR induced VM formation through TWIST1, we applied rescue experiments with add TWIST-cDNA (oeTWIST1) into SW839 cells and TWIST1-shRNA (shTWIST1) into 786O cells **(Fig. 2C-D)**. The results revealed that shTWIST1 in 786O cells could partially reverse the elevated VM formation induced by AR overexpression. In contrast, oeTWIST1 in SW839 cells increased VM formation, especially after knocking down the endogenous AR **(Fig. 2E, 2D assay and 2F, 3D assay)**. Similarly, Harmine (39, 40), which is a TWIST1 inhibitor causing its protein degradation **(Fig. 2G)**, could dramatically block the effect of overexpressing AR on VM formation of RCC cells **(Fig. 2H-I)**.

Together, results from **Fig. 2A-I** indicate that AR may function through modulating TWIST1 expression to influence the matrigel-coated 2D and collagen I based 3D VM formation *in vitro*.

### Human clinical sample analysis for TWIST1 expression in the ccRCC vs. para-tumor non-cancerous tissues

To examine the role of TWIST1 in ccRCC clinical samples, we detected its expression by IHC in 23 pairs of samples derived from ccRCC tumors and adjacent normal tissues **(Fig. 2J)** and the results showed a higher expression of TWIST1 was detected in tumors than in paired adjacent normal tissues (p < 0.001; n=23) **(Fig. 2K)**, which was consistent with the results extracted from TCGA mRNA database **(Fig. 2M)**. Furthermore, AR is positively correlated with TWIST1 based on 51 ccRCC IHC samples **(Fig. 2L)**. Additionally, based on TCGA online database UALCAN (http://ualcan.path.uab.edu/), we found elevating TWIST1 levels (using 25% of samples with the highest TWIST1 expression) led to a significantly lower survival rate in ccRCC patients **(Fig. 2N)**. Additionally, as shown in **SFig. 1B**, VM occurred more frequently in the TWIST1 high expression group.

### Mechanism dissection of how AR alters TWIST1 expression: *via* modulating the lncRNA TANAR expression

Since AR could elevate TWIST1 expression at both protein and mRNA levels **(See Fig. 2A-B)**, we then focused on whether AR increases TWIST1 expression *via* transcriptional regulation. Results from qPCR assay performed on 786O and SW839 cells treated for 2 hours with 10 nM DHT and 10 μM Enz respectively, showed no difference in mRNA levels, indicating AR may not regulate TWIST1 transcriptionally **(SFig. 2A)**. Intriguingly, after treating 786O oeAR cells or SW839 shAR cells with Actinomycin D for different periods of time to measure the decay of TWIST1 mRNA, we found that oeAR resulted in an increase in the half-life of TWIST1 mRNA, whereas shAR decreased its half-life implying that AR could stabilize TWIST1 mRNA. To further dissect the detailed mechanism, we tested whether AR could regulate miRNA targeting TWIST1 to exert its function **(Fig. 3A; B)**. As miRNA usually destabilizes its target genes in the Ago2-containing RISC complex (RNA-induced silencing complex) (41, 42), we performed an immunoprecipitation of Ago2 followed by detection of TWIST1 mRNA. The results showed that overexpressing AR contributed to a slight increase of TWIST1 mRNA level in the Ago 2 complex, indicating that AR may not modulate TWIST1 *via* altering miRNAs **(SFig. 2B)**.

**Fig. 3.** Mechanism dissection of how AR can increase TWIST1 expression: *via* lnc-TANAR. (A-B) 786O cells were transfected with vector control (pWPI) or AR-cDNA (oeAR) (A) and SW839 cells were infected with mock control (pLKO) or AR shRNA (shAR) (B). After 48 h, cells were incubated with 2 μg/mL actinomycin D for 0, 1, 2, 3, 4, and 5 hours. Total RNA was then analyzed by qRT-PCR to examine TWIST1 mRNA stability. (C) Bioinformatics analysis of potential lncRNAs that are associated with RCC resistance to Sunitinib in GSE69535 dataset and with predicted binding site to the mRNA of TWIST1 based on the RNA-RNA interaction prediction software, http://rtools.cbrc.jp/. (D) Real-time PCR of 19 potential candidate lncRNAs in SW839 cells infected with knocked down AR (shAR) compared with vector control (pLKO) and with adding AR-cDNA (oeAR) compared with scramble control (pWPI). (E) RNA pull-down assay *via* TWIST1 mRNA biotin identifies candidate lncRNAs ENST00000425110.1, and ENST00000377977.3, which can bind to TWIST1 mRNA more in 786O oeAR cells compared with 786O pWPI cells. (F) The qRT-PCR assay showed sh-ENST00000425110.1 and sh-ENST00000377977.3 could partly block oeAR increased TWIST1 mRNA level in 786O cells. (G-I) Knocking down TANAR reverses oeAR effect on TWIST1 protein level (G) and 2D (I, left) & 3D (I, right) formation in 786O cells. Quantitation of H in (I). (J-L) Overexpressing TANAR could partly reverse knock down AR effect on TWIST1 protein level (H) and 2D (K, left) & 3D (K, right) formation. Quantitation of K in (L). (M) Real time-PCR for TANAR, Malat1, and GAPDH from RNA extracted from nuclear and cytoplasmic fractions. (N) RNA fluorescence in situ hybridization (FISH) demonstrated that TANAR was localized in both cytoplasm and nucleus of 786O and SW839 cells.. Data are expressed as mean±S.D.. *p < 0.05, **p < 0.01, and ns=not significant, compared to the controls.

We then moved to the lncRNAs, as recent studies indicated that lncRNA may bind to the 5’UTR of target mRNA to regulate mRNA level (43, 44). To examine whether lncRNAs might be involved in regulating TWIST1 mRNA level, we first applied a bioinformatic analysis to identify potential lncRNAs capable of interacting with TWIST1 mRNA 5’UTR and overlapped the results with lncRNAs highly related to Sunitinib resistance in RCC from the GSE69535 dataset **(Fig. 3C)**. As shown in **Fig. 3D,** 5 candidates (ENST00000436510.1, ENST00000471626.1, ENST00000593604.1, ENST00000425110.1 and ENST00000377977.3) changed significantly after altering AR in 786O and SW839 cells. Furthermore, results from the RNA pull-down assay *via* TWIST1 mRNA showed that overexpressing AR in 786O cells could lead to ENST00000425110.1, and ENST00000377977.3 binding more to TWIST1 mRNA compared with pWPI group **(Fig. 3E)**.

At the same time, knocking down these candidates **(SFig. 2C)** indicated that knockdown of ENST00000425110.1 and ENST00000377977.3 could suppress TWIST1 expression more in the oeAR group than in the control group **(Fig. 3F)**. In addition, western blot assays revealed that only ENST00000425110.1 overexpression could rescue the decline in TWIST1 protein induced by knocking down AR. As expected, the overexpressed AR-elevated TWIST 1 protein level decreased drastically *via* knockdown of ENST00000425110.1 **(Fig. 3G and H; SFig. 2D-F)**. Consistent with these results, we also found that overexpressing ENST00000425110.1 in SW839 cells could increase VM formation, especially after knocking down the endogenous AR **(Fig. 3K)**. In contrast, overexpressed AR-increased ccRCC VM could be partially blocked by adding ENST00000425110.1-shRNA in 786O cells **(Fig. 3I-J and K-L)**. Therefore, we named this lncRNA as TANAR (Twist1 Associated long Noncoding RNA regulated by AR) based on the above analyses and focused on this lncRNA for the remaining studies.

Through Ensemble software and Lncpedia database, we found that the position of TANAR, with little protein-coding function, is in Chromosome 2: 240,981,515-240,986,072 **(SFig. 3A-B)**. Furthermore, subcellular fraction analysis, as well as FISH assay, revealed that TANAR was distributed in both cytoplasm and nuclei of cells **(Fig. 3M-N)**, consistent with online analysis (http://www.csbio.sjtu.edu.cn/bioinf/lncLocator/) predicting the location of TANAR **(SFig. 3C)**.

Together, from **Fig. 3A-N; SFig. 2A-F; and SFig. 3A-C**, we found a novel lncRNA-TANAR that is regulated by AR *via* altering TWIST1 expression and inducing VM formation in ccRCC cells.

### Mechanism dissection of how AR alters TANAR expression: *via* transcriptional regulation

To further dissect the potential molecular mechanism how AR regulates the TANAR expression at the transcriptional level, we applied the Ensembl and PROMO 3.0 websites to search for the androgen-response-elements (AREs) in the 2 kb region of the TANAR promoter by using JASPAR database **(Fig. 4a)**, and detected five putative AREs (I −1150nt to −1136nt, II/III −602nt to −586nt and VI/V −94nt to −41nt) **(Fig. 4b)**. Next, we performed the chromatin immunoprecipitation (ChIP) assay, and results revealed that AR could specifically bind to ARE II&III, but not the other AREs **(Fig. 4c)**. Furthermore, we mutated the critical sequences of ARE II/III and inserted the mutant (MT) promoter region of TANAR into pGL3 luciferase plasmid as well as the wild-type (WT) promoter **(Fig. 4D)**. As expected, the luciferase assay results showed that knocking down AR significantly lessened luciferase activity in SW839 cells transfected with WT reporter, but not in the cells with the MT reporter **(Fig. 4E left).** In contrast, overexpressing AR could drastically increase luciferase activity in 786O cells with the WT reporter, but not in the cells with the reporter containing the MT ARE **(Fig. 4E right)**.

**Fig. 4.**
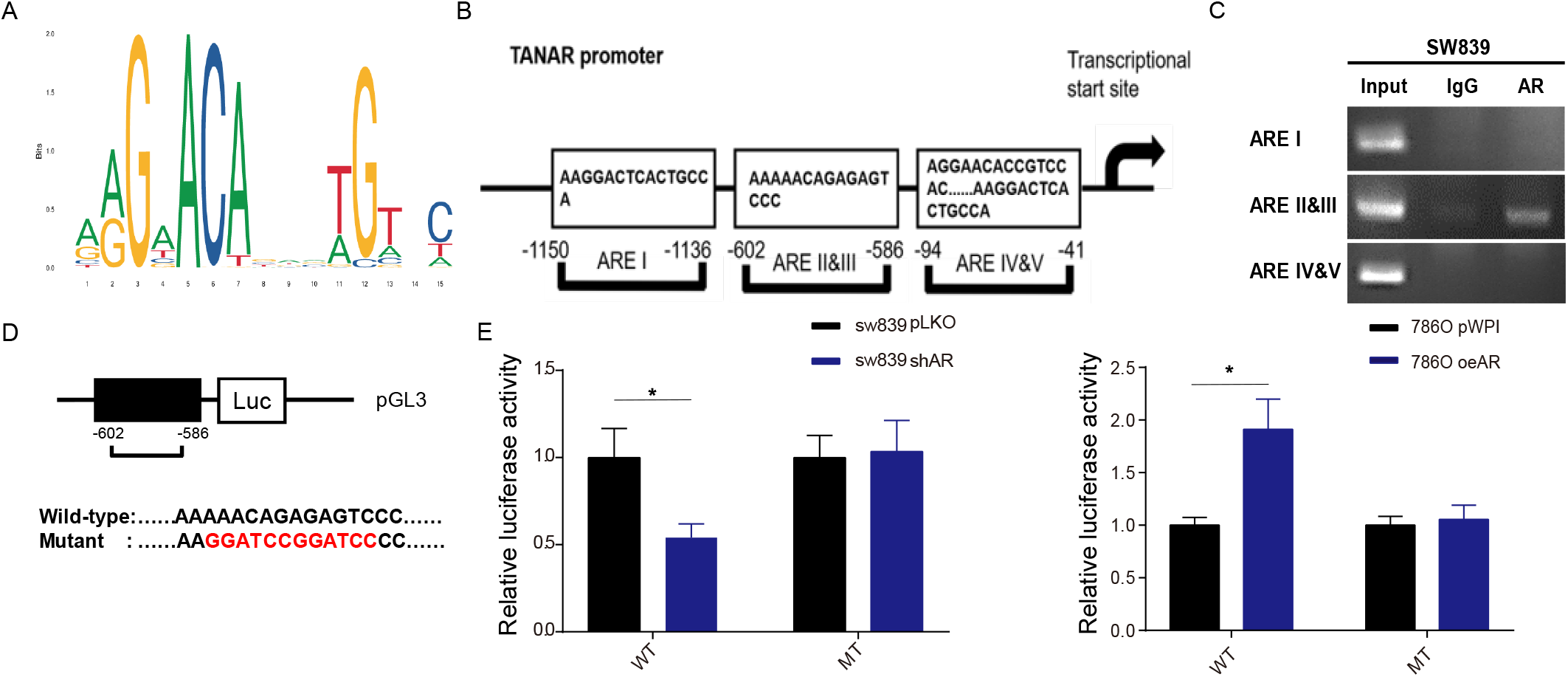
Mechanism dissection how AR regulates TANAR expression: *via* transcriptional regulation. (A)ARE motif sequence. (B) The structure of ARE binding site in 2kb TANAR promoter region. (C) CHIP assay confirmed AR could directly bind with TANAR AREII/III (−602nt to −586nt). (D) schematic diagram of wild type and mutant pGL3-TANAR promoter-reporter constructs. (E-F) Co-transfection of ARE wild type (WT) or Mutant (MT) TANAR promoter pGL3-Luciferase plasmids into SW839 cells with pLKO or shAR (E) and 786O cells with pWPI or oeAR (F).The luciferase reporter assay was performed to detect promoter activity. The data are the means ± S.D. *p < 0.05, and ns=not significant, compared with control.

Together, results from **Fig. 4A-E** suggested that AR could directly increase TANAR expression transcriptionally through binding to the ARE II/III.

### Mechanism dissection of how TANAR can alter TWIST1 expression: *via* modulating nonsense-mediated mRNA decay by competitive binding of UPF1 to TWIST1 mRNA

To dissect the molecular mechanism underlying AR/TANAR’s regulation on TWIST1 expression, especially the impact of AR on TWIST1 mRNA stability (**See Fig. 3A, B**), we hypothesized that TANAR could modulate TWIST1 mRNA stability *via* the TANAR-TWIST1 interaction, as was shown with the presence of TANAR in TWIST1 pull-down assay **(See Fig. 3E)** (43, 45, 46). Indeed, mRNA stability assay, using treatment with Actinomycin, revealed that knocking down TANAR could block the overexpressing AR-increased TWIST1 mRNA stability **(Fig. 5A)**. Furthermore, to test whether this binding is crucial for TANAR’s regulation on TWIST1 mRNA, we constructed WT TANAR cDNA sequence and mutated it by deleting the TANAR-TWIST presumptive binding region **(Fig. 5B; SFig. 4A; SFig. 4B)**. Compared to the pWPI group, 786O cells transfected with MT TANAR-cDNA failed to increase TWIST1 mRNA and protein level **(Fig. 5C-D)**. Moreover, mRNA stability assay, using Actinomycin treatment, showed that AR knockdown-decreased TWIST1 mRNA stability could be rescued only by adding wild type TANAR **(Fig. 5E)**. The FISH assay in SW839 cells also showed that TANAR could colocalize with TWIST1 mRNA (left panel), and we found that using Image J colocalization analyser the Pearson’s correlation coefficient (PCC) is 0.6554 (right panel) **(Fig. 5F).** Taken together, these data confirmed that the stabilizing effect of TANAR relies on its direct binding of TWIST1 mRNA.

**Fig. 5.**
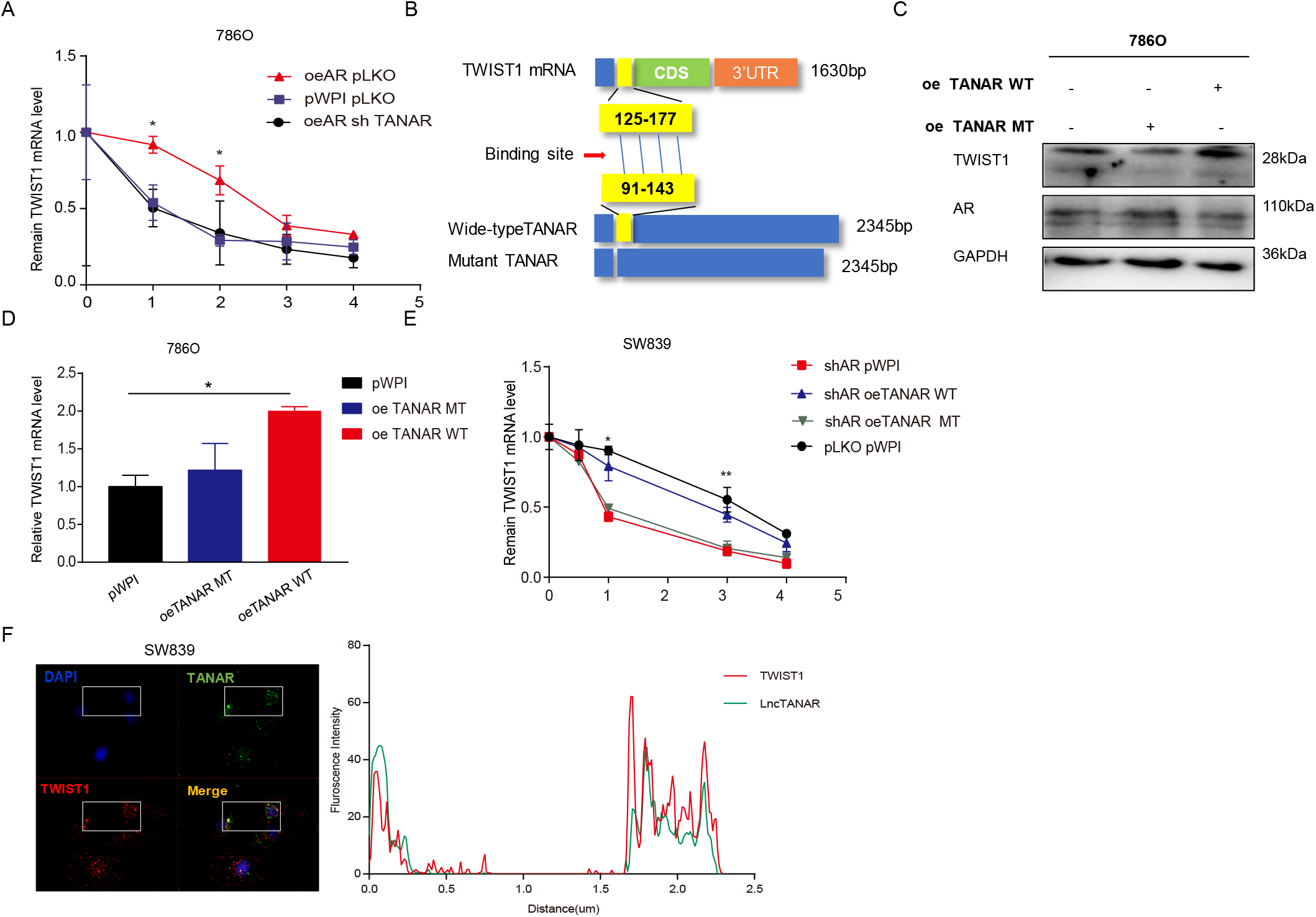
Mechanism dissection how TANAR modulates TWIST1 expression: *via* stabilizing TWIST1 mRNA. (A) 786O cells transfected with oeAR pLKO, pWPI pLKO, or oeAR shTANAR were treated with 2 μg/ml actinomycin D for designed times. Real-time RT-PCR was performed to examine the mRNA stability of TWIST1. (B) Sketch map showed construct of wild-type TANAR and mutant TANAR. (C) Western blot assay showed the influence of ectopic wild-type (WT) and mutant (MT) TANAR on TWIST1 protein levels in 786O cells. (D) RT-PCR showed the influence of ectopic WT and Mut TANAR on TWIST1 mRNA level in 786O cells. (E) SW839 cells expressing shAR pWPI, shAR oeTANAR WT, shAR oeTANAR MT or pLKO pWPI, were treated with 2 μg/ml actinomycin D for the designed periods of time. Real-time RT-PCR was performed to examine the mRNA stability of TWIST1. (F) Co-localization of TANAR (green, upper right) and TWIST1 mRNA (red, lower left) signals using RNA FISH. Nuclei were stained with DAPI (upper left lane). The lower right frame indicates the co-localized region as calculated by Image J colocalization analyser software. The data are the means ± S.D. *p < 0.05, **p < 0.01, and ns=not significant, compared with control.

Intriguingly, based on Ensembl database (http://uswest.ensembl.org/) and individual-nucleotide-resolution UV cross-linking and immunoprecipitation (iCLIP) performed by Zünd David (47), TWIST1 mRNA could interact with the indispensable component for nonsense-mediated mRNA decay (NMD)-UPF1 through at least 12 binding sites **(S. Table1)**, and then be regulated by NMD process *via* rapid degradation as its mRNA conforms to a canonical NMD structure **(SFig. 4C)** (27).

To test whether TANAR could alter TWIST1 mRNA *via* influencing NMD, we performed western blot to see the effect of TANAR on NMD core factors, UPF1 and SMG1 (48) expression. We found that overexpressing TANAR could not reduce either UPF1 or SMG1 protein level **(SFig. 4D)**. However, the RNA immunoprecipitation assay by pull-down of TWIST1 mRNA using biotin-conjugated antisense oligonucleotide revealed that ectopic expression of wild-type TANAR could effectively diminish TWIST1 mRNA-UPF1 interaction in ccRCC, while their interaction was increased after knocking down TANAR **(Fig. 6A-D)**. Consistent with that, overexpressing wild-type TANAR, but not mutant TANAR, reduced TWIST1 level in the UPF1 immunoprecipitation **(Fig. 6E)**. Knocking down TANAR significantly increases the TWIST1 level in UPF1 immunoprecipitation **(Fig. 6F)**.

**Fig. 6.**
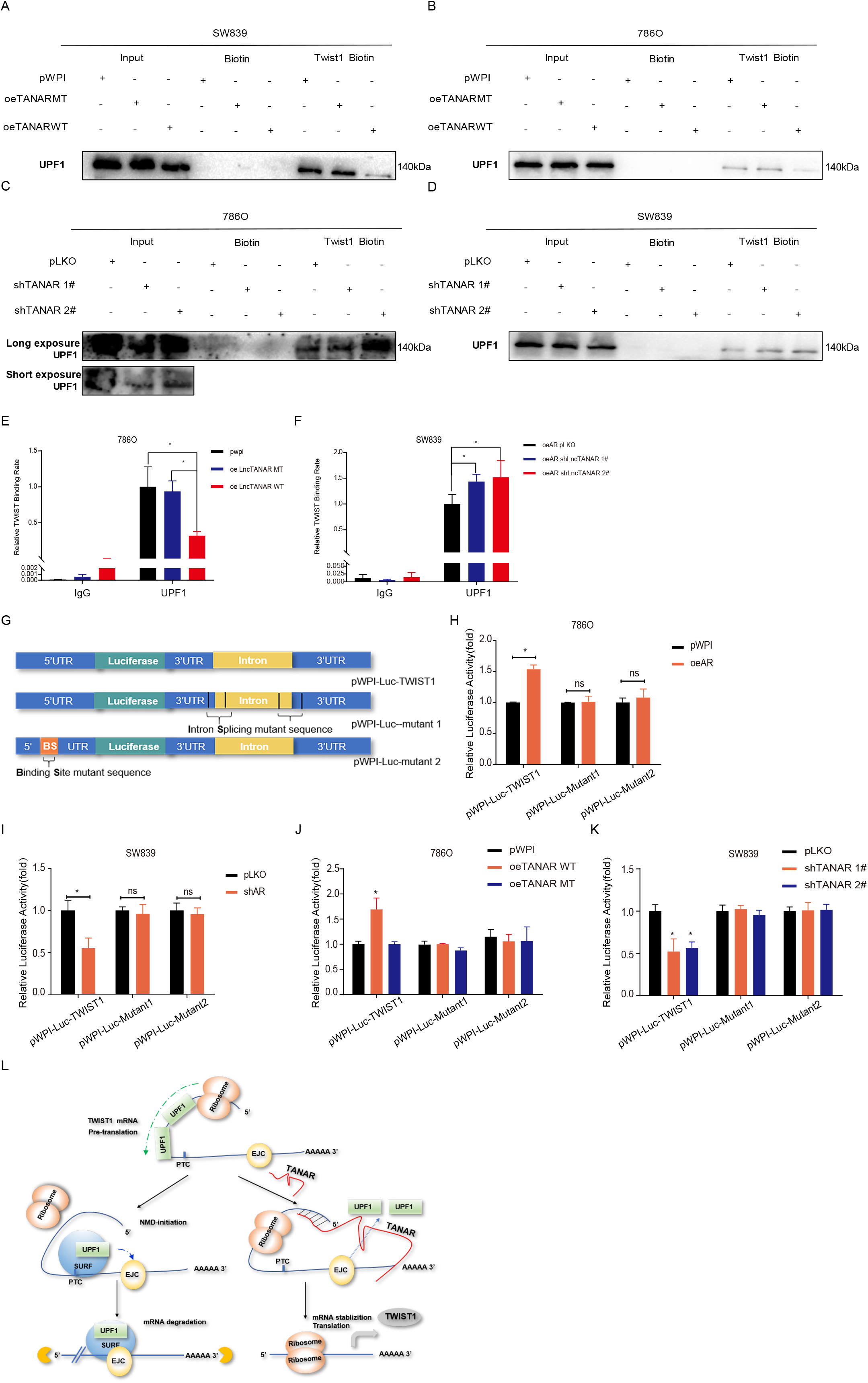
Mechanism dissection how TANAR modulates TWIST1 stability: *via* diminishing TWIST1 nonsense-mediated decay by interrupting the interaction between UPF1 protein and TWIST1 mRNA. (A-D) SW839 (A and D) or 786O (B and C) cells were transfected with pWPI, oeTANAR MT, oe TANAR WT, shTANAR 1# or shTANAR 2# as indicated. Cell lysates were incubated with *in vitro* biotin-labeled sense or antisense probes against TWIST1 mRNA for the RNA pull-down assay. Western blot assays were performed to test levels of UPF1 in sediments from the pull-down. (E-F) Cell lysates of 786O (E) cells expressing control pWPI, oe TANAR MT, or oe TANAR WT and SW839 (F) cells expressing oeAR control pLKO, oeAR shTANAR 1#, or oeAR shTANAR 2# were incubated with biotin-labeled sense or antisense probes against TWIST1 mRNA for RNA pull-down assay and RT-PCR analysis to test TWIST1 mRNA levels. (G) Sketch map showed the construct of pWPI-luc-TWIST1 and pWPI-luc-mutant 1 (intron splicing mutant) and pWPI-luc-mutant 2 (binding site mutant). (H-I) Co-transfection of 786O cells with pWPI or oeAR (H) or SW839 cells (I) with pLKO or shAR and both cells types with pWPI-luc-TWIST1, pWPI-luc-mutant 1 or pWPI-luc-mutant 2 plasmids. The luciferase reporter assay was performed to detect promoter activity. (J-K) Co-transfection of pWPI-luc-TWIST1, pWPI-luc-mutant1 or pWPI-luc-mutant2 plasmids into SW839 cells with pLKO or shTANAR1# &2# (K), and 786 cells with pWPI, oeTANAR WT or oeTANAR MT (J). The luciferase reporter assay was performed to detect promoter activity. (L) Schematic diagram: Lnc TANAR could directly bind to TWIST1 mRNA, diminishing its interaction with NMD core factor, UPF1, thus stabilize TWIST1 mRNA. **Abbreviations**: EJC: exon junction complex; PTC: premature termination codon; SURF:SMG1-UPF1-eRF1-eRF3 complex. The data are the means ± S.D.. *p < 0.05, **p < 0.01, and ns=not significant, compared with control.

In order to further validate the function of TANAR on the NMD process of TWIST1, we generated luciferase report constructs replacing TWIST1 coding region with luciferase cDNA while maintaining the overall gene structure of TWIST1 genomic locus (pWPI-luc-TWIST1). We also generated two mutant reporters with mutations in the conserved splicing sequence to eliminate the NMD initiation (pWPI-luc-mutant 1) and TANAR-interacting region in the 5’UTR (pWPI-luc-mutant 2) **(Fig. 6G)**. As expected, the results showed that overexpressing AR could significantly increase luciferase activity in 786O cells with pWPI-luc-TWIST1, but not in the cells with either pWPI-luc-mutant 1 or pWPI-luc-mutant 2 **(Fig. 6H)**. Consistent with this, knocking down AR significantly reduced luciferase activity in SW839 cells transfected with pWPI-luc-TWIST1, but not in the cells with either of the 2 mutant plasmids **(Fig. 6I)**. When treating cells with DHT or Enz, we found the same tendency **(SFig. 4E-F)**. Moreover, only 786O cells cotransfected with WT TANAR cDNA sequence and pWPI-luc-twist1, not with TANAR MT TANAR-cDNA or pWPI and pWPI-luc-mutant 1 or pWPI-luc-mutant 2, showed increased luciferase activity **(Fig. 6J)**. The luciferase activity decreased only when knocking down TANAR in SW839 cells transfected with pWPI-luc-TWIST1 **(Fig. 6K)**.

Together, results from **Fig. 6A-K and SFig. 4A-F** suggest TANAR suppresses UPF1-TWIST1 mRNA interaction, thus reduces nonsense-mediated decay of TWIST1 mRNA to increase its mRNA stability **(Diagram in Fig. 6L)**.

### Preclinical study using *in vivo* mouse model to confirm the role of AR/TANAR/TWIST1 axis in ccRCC VM

To further test the validity of the above *in vitro* data, we applied the orthotopic ccRCC xenograft mouse model. We generated stable clones of 786O cells with luciferase expression with overexpressed AR and/or sh-TANAR as well as a control, with 5 mice/group (1: Scr +luc; 2: oeAR + luc; 3: sh-TANAR + luc; and 4: oeAR + sh-TANAR + luc). A total of 1 × 10^6^ 786O cells mixed with matrigel were inoculated into the left kidney capsule of nude mice and tumor progressions were evaluated *via* the non-invasive *in vivo* imaging system (IVIS). After 6 weeks, we observed that increased bioluminescence signals in the whole mice **(Fig. 7A-D)**, as well as in the left kidney in the oeAR xenografted groups while knocking down TANAR could partly reverse the oeAR-increased high radiance in ccRCC xenografts. Furthermore IVIS showed a dramatic increase of metastatic luciferase signals (as seen in the liver, intestine, diaphragm, spleen, and testis) in the oeAR group and less metastasis occurred in the sh-TANAR group than control group **(Fig. 7E-J)**. Upon animal sacrifice, and retrieval of tumors and metastases, we found targeting the TANAR with sh-lncRNA-TANAR could partly reverse the oeAR-promoted ccRCC metastasis foci **(Fig. 7K)**. Importantly, results from IHC staining demonstrated that oeAR led to increase the TWIST1 expression as well as VM vessel area (PAS+/CD31-), which could be partly reversed by sh-TANAR **(Fig. 7L-N)**.

**Fig. 7.**
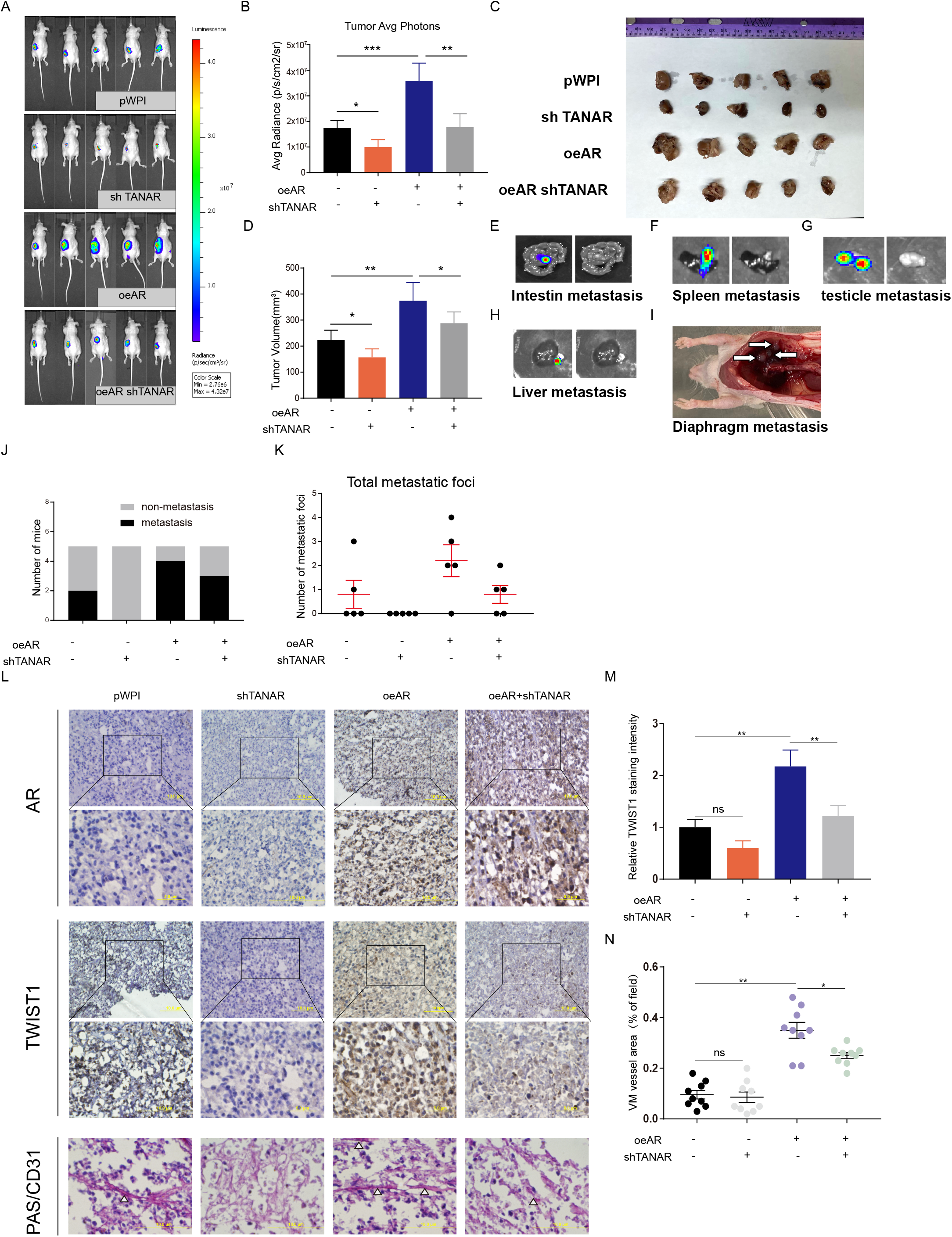
Preclinical study using in vivo mouse model to test the role of AR and TANAR in ccRCC VM formation. (A) IVIS images of mice harboring RCC tumors after orthotopically implanting 786O pWPI,786O shTANAR,786O oeAR and 786O oeAR-shTANAR cells into nude mice (N=5) for 6 weeks.(B) Tumor average photons for ccRCC from xenograft mice described above. (C) Images of tumors are presented after mice were sacrificed and the tumor volume in each group were observed and measured (D). (E-I) Representative organ bioluminescent images showing metastasis from testicles, liver, intestine, spleen, and diaphragm metastasis. (J) Quantification of the metastasis in the four groups of mice. (K) Quantification of the total metastatic foci. (L) Representative images of IHC staining for AR, TWIST1, and VM vessel area (White triangles show PAS+/CD31-tumor cell-dependent vessels) in mice. (M) Quantification of relative IHC staining intensity for TWIST1 expression. (N) Evaluation of area percentage of VM vessels. The data are the means ± S.D. *p < 0.05, **p < 0.01, ***p < 0.05, and ns=not significant, compared with control.

Together, results shown in **Fig. 7A-N** are consistent with the *in vitro* cell lines studies and proved that targeting AR/lncTANAR/TWIST1 axis could prevent VM formation in the tumor tissue and consequently suppress tumor progression and metastasis.

## Discussion

The epithelium-dependent angiogenesis has been long regarded as the sole blood supplement initiated by proangiogenic factors. Due to the highly vascularized characteristic of ccRCC, hematogenous metastasis ultimately occurred in most late-stage patients with ccRCC. The anti-angiogenesis medicines, RTKIs, targeting VEGF and its receptor, such as sunitinib, remain the first-line treatment with median survivals up to 30 months (49, 50). However, acquired drug resistance will inevitably occur in most metastatic ccRCC patients within 12 months (51). The conception of VM, epithelium-independent angiogenesis, <u>reinforces</u> the tumor neovascularization theory, indicating that targeting critical factors regulating VM formation in the tumor could be a novel anti-angiogenesis therapy to stop tumor cells (36).

Recent literature revealed that VM formation, recognized as PAS-positive and CD31 negative, is an independent and relevant prognostic criterion for disease-free survival in patients with ccRCC (52, 53). Similarly, based on 51 ccRCC clinical samples, we found a higher stage tends to possess more VM channels (Fig. 1B), partly explaining why VM presence leads to poor prognosis in ccRCC.

Interestingly, across our cohort, VM formation exists predominantly in males compared to females in Stage II&III (Fig. 1C). However, consistent with a previous study (53), once we brought in Stage I clinical data, VM formation shows no significant difference between male and female (SFig.1A), which could be partly due to scarce incidence of VM in Stage I patients in a larger male patient population lowering overall VM and a lack of gender difference (Fig.1B).

Mounting evidence has demonstrated that AR functions as an oncogene in ccRCC, promoting progression and hematogenous metastasis (5, 6, 23), despite the presence of few retrospective literatures based on TCGA database implied AR may contribute to better prognosis in RCC (9). However, its role in VM formation remains unclear. Based on our Formalin-fixed, Paraffin-embedded (FFPE) ccRCC tissue samples, expression of AR was detected more frequently in patients with higher VM presence, indicating that AR was positively correlated with VM formation (R=0.3947; P < 0.001; Fig. 1D-E). Although AR could act as a marker for the aggressive VM, the clinical data revealed that some AR negative and VM positive cases do exist, suggesting that AR may not be the exclusive regulator of VM. Consistent with pathological data, ectopic expression or DHT-induced activation of AR increases, whereas knock down of AR or Enz-induced inactivation of AR decreases, the classical Matrigel-coated 2D VM formation. In 2017, D. O. Velez developed a de novo 3D *in vitro* collagen-induced migration procedure to detect VM, which is broadly related to metastasis in solid human tumors including kidney cancer. Compared to the classical Matrigel-coated VM formation culture system, a pseudo 3D architecture where cancer cells are embedded in the extracellular matrix (ECM) contacting with cover glass, this novel Collagen I-based 3D culture system prevents cancer cells from the influence of the coverslip, thus a more physiologically relevant *in vitro* VM model (19). We utilized the 3D Collagen-I-induced model to further confirm AR influence on VM formation. Moreover, we have also proved the role of AR in the formation of VM *in vivo via* the orthotopic implantation model, finding increased VM formation, increased tumor progression and more metastases in xenografts established by cells with AR overexpression. Taken together, our study highlights the significance of AR in both *in vitro* and *in vivo* VM formation in a physiological way for the first time.

We found that AR can increase ccRCC VM formation *via* up-regulating TWIST1. TWIST1 is a basic helix-loop-helix transcriptional factor, which plays important roles in epithelial-mesenchymal transition and VM in diverse types of tumors (39, 54, 55). Previous literature based on 163 ccRCC clinical samples revealed that elevated levels of TWIST1, which is mainly localized in the cytoplasm of ccRCC cells (98.8%), was closely associated with higher stage, vascular invasion, and poor prognosis in RCC (56). In addition, it has been reported that TWIST1 could transcriptionally regulate VE-cadherin, a transmembrane protein responsible for cell-cell adhesion and VM formation, in multiple types of tumor cells (57, 58). Not surprisingly, our clinical tissue samples demonstrated that TWIST1 expression is positively correlated with VM formation (SFig. 1C; R=0.34; P < 0.001), which is consistent with our *in vivo* or *in vi*tro data. Furthermore, we treated ccRCC cell with Harmine, a naturally occurring beta-carboline alkaloid widely used herb, as a TWIST1 inhibitor selected through the unbiased screen and validated in the lung cells (39, 40), and found that it led to a dramatic degradation of TWIST1 protein at either 10 μM or 40 μM while also drastically diminished VM formation in ccRCC cells. Thus, targeting TWIST1 signaling might be a promising therapy for ccRCC with high VM formation.

Increasing evidence indicates that TWIST1 could be post-transcriptionally modulated *via* noncoding RNA (59–61). The lncRNAs could sponge microRNAs to regulate of the TWIST1 expression, and then influence tumorigenesis. Moreover, NMD, which is a conserved cellular mRNA surveillance system, can degrade TWIST1 transcripts *via* recognition of Premature termination codons in TWIST1 mRNA (62, 63). On the one hand, cancer cells have utilized NMD to decrease gene expression by deliberate reduction of vital tumor-suppressor mRNAs (64). On the other hand, cancer cells could fine-tune NMD activity to adapt to harsh microenvironments (65). However, whether lncRNA is involved in the regulation of mRNA stability *via* NMD still remains not clear. In this study, we identified a novel lncRNA-TANAR, as an AR-transcriptionally-regulated lncRNA, is able to promote TWIST1 mRNA stability by suppressing NMD *via* competitive binding with the Up-frameshift protein 1 (UPF1) to the TWIST1 mRNA 5’UTR. Intriguingly, previous studies showed that UPF1 mostly targets 3’ untranslated region of mRNA to exert NMD function (66) while TANAR binds to the 5’UTR region of the TWIST1 mRNA. To further validate the role of NMD in regulating TWIST1 mRNA, we scrutinized iCLIP-seq data and found 12 confirmed UPF1 protein-TWIST1 mRNA binding sites (27), partly overlapping with the predicted TANAR-TWIST1 mRNA interaction region. Based on our RIP assays and luciferase analyses (Fig. 6A-K), we show lncRNA-TANAR increases TWIST1 mRNA stability *via* directly binding to its 5’UTR with disruption of UPF1 initiating nonsense-mediate TWIST1 mRNA decay.

This suggests a role of 5’UTR in regulating NMD likely through the loop formation between the 5’UTR and 3’UTR of mRNA.

In conclusion, we characterized a new long noncoding RNA TANAR transcriptionally regulated by AR, which influences VM formation by decreasing TWIST1 mRNA nonsense-mediated decay. Targeting the AR/lncRNA-TANAR/TWIST1 axis could be a promising strategy for the development of a better treatment of ccRCC.

## Acknowledgments

This work was supported by George Whipple Professorship. We thank Karen Wolf for help preparing the manuscript.

## Ethics approval and consent to participate

Ethical consent was approved by the Committees for Ethical Review of Research involving Human Subjects at Harbin Medical University. Written informed consent was obtained from each patient prior to sample collection. The animal experiments were approved by the Use Committee for Animal Care at University of Rochester Medical center.

## Consent for publication

Not applicable.

## Availability of data and materials

The datasets used and analyzed during the current study are available from the corresponding author on reasonable request.

**SFig. 1.** (A) VM channels per high-quality frame (HPF) in Male (n=36) and Female (n=15) patients samples. (B) VM formation was quantified in TWIST1-high (n = 24) and TWIST1-low (n = 27) tumor tissues. Red=VM Negative and black=VM Positive.

**SFig. 2** (A) RT-PCR analyses were performed after treated 786O (left) and SW839 (right) cells with Ethanol/10 nM DHT or DMSO/10 μM Enz, respectively. (B) Ago2 pull-down assays were performed for RT-PCR. The TWIST1 mRNA levels were detected in 786O cells transfected with pWPI or oeAR. (C) The RT-PCR was applied to validate the knockdown efficiency of 5 lncRNA candidates. (D) The RT-PCR was applied to validate the overexpression efficiency of ENST 00000425110.1 and ENST 00000377977.1. (E-F) Western blot assays were performed on 786O cells (E) transfected with pWPI+pLKO,pWPI+sh ENST00000377977,oeAR+pLKO and oeAR+sh ENST00000377977 and on SW839 (F) cells transfected with pLKO+pWPI, pLKO+oe ENST00000377977, shAR+pWPI and shAR+oe ENST00000377977.

**SFig. 3** (A) The chromosome location of TANAR based on ensemble software. (B) The protein-encoding possibility of lncRNA-TANAR predicted by LNCipedia.org. (C) The lncRNA-TANAR localization predicted by LncLocator.

**SFig. 4** (A) The potential binding site structure and joint secondary structure between TANAR and TWIST1 analyzed by online RNA-RNA predicting software (rtools.cbrc.jp). (B) The overexpression efficiency of TANAR wild type (WT) and mutant (MT) through RT-PCR. (C) Ensembl online database demonstrated the nonsense-mediated mRNA decay occurred in TWIST1 transcripts. (D) Western blot was performed to detect the TANAR impact on UPF1 protein expression. (E-F) Transfection of pWPI-luc-TWIST1, pWPI-luc-mutant 1 or pWPI-luc-mutant 2 plasmids into 786O cells with 10 nM DHT for 24 h (E) or into SW839 cells treated with 10 μM Enz for 24 h (F). The luciferase reporter assay was performed to detect promoter activity.. The data are the means ± S.D. *p < 0.05, and ns=not significant, compared with control.

